# Enhanced virus detection and metagenomic sequencing in patients with meningitis and encephalitis

**DOI:** 10.1101/2020.11.25.399394

**Authors:** Anne Piantadosi, Shibani S. Mukerji, Simon Ye, Michael J. Leone, Lisa M. Freimark, Daniel Park, Gordon Adams, Jacob Lemieux, Sanjat Kanjilal, Isaac H. Solomon, Asim A. Ahmed, Robert Goldstein, Vijay Ganesh, Bridget Ostrem, Kaelyn C. Cummins, Jesse M. Thon, Cormac M. Kinsella, Eric Rosenberg, Matthew P. Frosch, Marcia B. Goldberg, Tracey A. Cho, Pardis Sabeti

## Abstract

Meningitis and encephalitis are leading causes of central nervous system (CNS) disease and often result in severe neurological compromise or death. Traditional diagnostic workflows largely rely on pathogen-specific diagnostic tests, sometimes over days to weeks. Metagenomic next-generation sequencing (mNGS) is a high-throughput platform that profiles all nucleic acid in a sample. We prospectively enrolled 68 patients from New England with known or suspected CNS infection and performed mNGS from both RNA and DNA to identify potential pathogens. Using a computational metagenomic classification pipeline based on KrakenUniq and BLAST, we detected pathogen nucleic acid in cerebrospinal fluid (CSF) from 22 subjects. This included some pathogens traditionally diagnosed by serology or not typically identified in CSF, including three transmitted by *Ixodes scapularis* ticks (Powassan virus, *Borrelia burgdorferi*, *Anaplasma phagocytophilum*). Among 24 subjects with no clinical diagnosis, we detected enterovirus in two subjects and Epstein Barr virus in one subject. We also evaluated two methods to enhance detection of viral nucleic acid, hybrid capture and methylated DNA depletion. Hybrid capture nearly universally increased viral read recovery. Although results for methylated DNA depletion were mixed, it allowed detection of varicella zoster virus DNA in two samples that were negative by standard mNGS. Overall, mNGS is a promising approach that can test for multiple pathogens simultaneously, with similar efficacy to pathogen-specific tests, and can uncover geographically relevant infectious CNS disease, such as tick-borne infections in New England. With further laboratory and computational enhancements, mNGS may become a mainstay of workup for encephalitis and meningitis.

**Importance:** Meningitis and encephalitis are leading global causes of central nervous system (CNS) disability and mortality. Current diagnostic workflows remain inefficient, requiring costly pathogen-specific assays and sometimes invasive surgical procedures. Despite intensive diagnostic efforts, 40-60% of people with meningitis or encephalitis have no clear cause of their CNS disease identified. As diagnostic uncertainty often leads to costly inappropriate therapies, the need for novel pathogen detection methods is paramount. Metagenomic next-generation sequencing (mNGS) offers the unique opportunity to circumvent these challenges using unbiased laboratory and computational methods. Here, we performed comprehensive mNGS from 68 patients with suspected CNS infection, and define enhanced methods to improve the detection of CNS pathogens, including those not traditionally identified in the CNS by nucleic acid detection. Overall, our work helps elucidate how mNGS can become a mainstay in the diagnostic toolkit for CNS infections.

## Introduction

Meningitis and encephalitis are leading causes of central nervous system (CNS) disease, ranked as the 4th leading contributor to global neurological disability-adjusted life-years (1), often resulting in severe neurological compromise or death (2, 3). Traditional diagnostic workflows remain inefficient, requiring costly pathogen-specific diagnostics, serial cerebral spinal fluid (CSF) testing, and sometimes invasive surgical procedures. Despite these intensive diagnostic efforts, 40-60% of subjects with meningitis or encephalitis have no clear cause identified (2, 4–6).

Metagenomic next-generation sequencing (mNGS) offers a unique opportunity to circumvent some of these challenges. mNGS consists of unbiased sequencing of all nucleic acid in a sample and computational classification of reads to identify potential pathogens (7–9). This technique successfully detected a range of pathogens, including bacteria (10–12), fungi (13), protozoa (14), and viruses (15–17) in subjects with CNS infection. mNGS is increasingly used as a clinical diagnostic test (18–20), and criteria for test performance have been described, though not yet standardized (21–23).

In this study, we prospectively enrolled 68 patients with known or suspected CNS infection and performed mNGS from both RNA and DNA to identify pathogens. We focused laboratory and analysis methods on viral nucleic acid detection since viruses are the most common type of pathogen detected in CNS infection (4, 5, 24, 25). Goals for this study were to assess the utility of standard mNGS in identifying CNS pathogens and examine enhanced laboratory techniques for improving analytic sensitivity, including hybrid capture (HC) of viral nucleic acid and methylated DNA depletion (MDD).

## Methods

### Subject enrollment and clinical characterization

The Prospective Encephalitis and Meningitis Study (PEMS) is a prospective cohort study enrolling adults who presented to Massachusetts General Hospital (MGH) with confirmed or suspected CNS infection. A total of 136 subjects enrolled in the PEMS between April 2016 and December 2017, of whom 122 had available CSF. Immunocompetent patients with CSF white blood cell count (WBC) < 5 cells/ul (n = 40) were excluded as unlikely to have infectious meningitis or encephalitis. Additional exclusions included encephalitis due to nosocomial bacteria, or bacteria and fungi that would be challenging to distinguish from common laboratory contamination in mNGS (n = 14) (Supplementary Table 1, DOI: 10.6084/m9.figshare.13266506). Sixty-eight subjects were included in mNGS analysis (Figure 1). This study was approved by the Partners Institutional Review Board under protocol 2015P001388. Further details are in Supplementary Methods.

**Figure 1.**
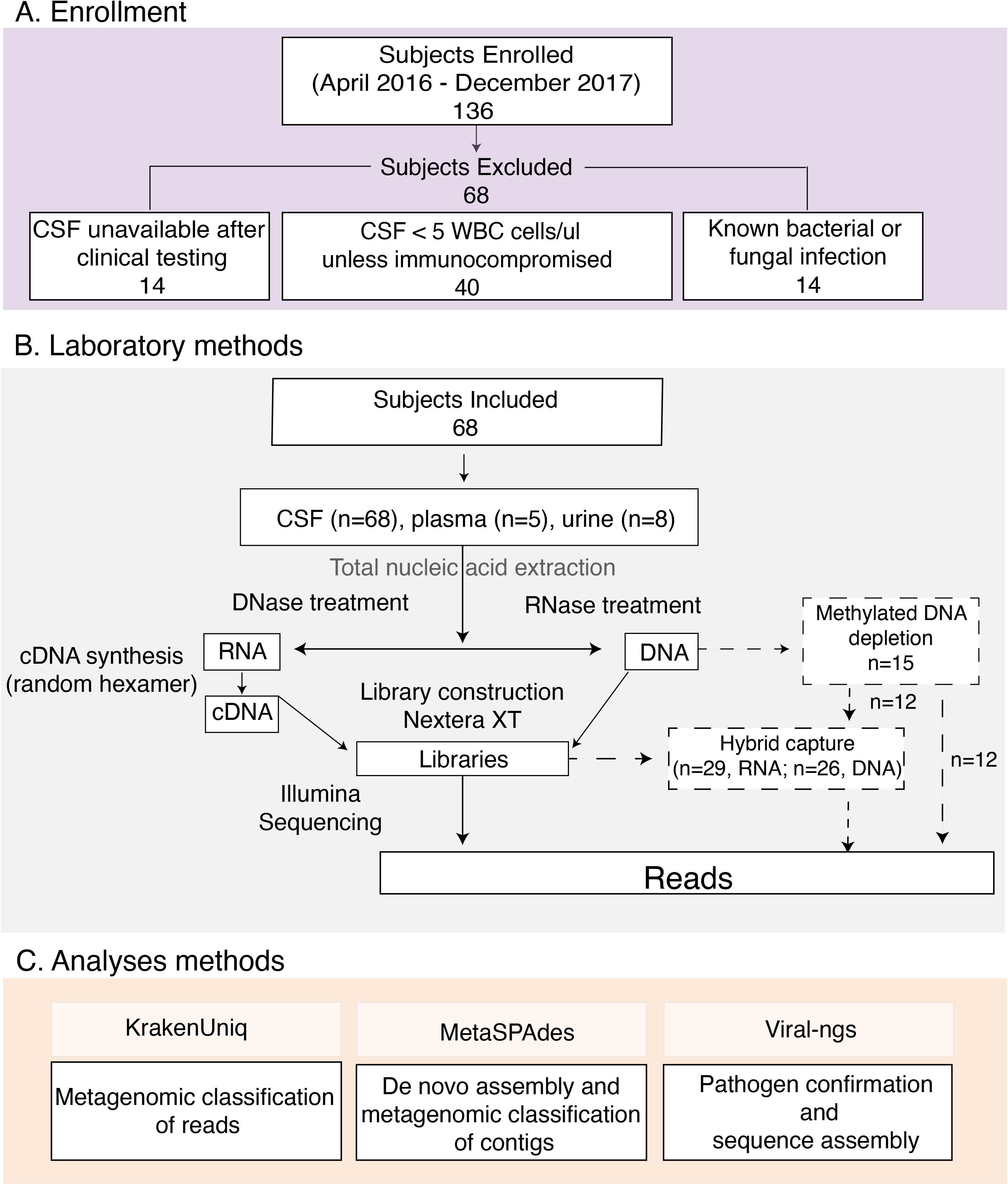
Overview of methods for subject selection and metagenomic next generation sequencing. Enrollment (A), laboratory methods (B), and analysis methods (C) are shown. Enhanced laboratory methods for methylated DNA depletion and hybrid capture (dashed lines) were included for a subset of the samples as shown. Abbreviations: CSF, cerebrospinal fluid; WBC, white blood cells; cDNA, complementary DNA; μl, microliter

### Nucleic acid isolation and standard mNGS

To minimize environmental contamination from viruses studied in the research laboratory, nucleic acid extraction and library construction were performed in an isolated workspace with limited access, extensive decontamination, and strict oversight of supplies, storage areas, and reagents. As a negative control, water and/or CSF from an uninfected patient (Negative CSF) was included with each batch starting from nucleic acid isolation. Nucleic acid was extracted from 140μl of CSF, urine, or plasma stabilized with linear acrylamide using the QIAmp Viral RNA Mini Kit (Qiagen). The elution was split into two fractions for RNA and DNA sequencing, and External RNA controls consortium (ERCC) spike-in oligonucleotides were added to each. Methods for cDNA synthesis have been previously described (26, 27). Both DNA and RNA libraries underwent tagmentation with the Nextera XT DNA Library Prep Kit (Illumina), and were pooled and sequenced on HiSeq and MiSeq machines using paired-end 100 or 150bp reads. Methods are outlined in Figure 1B, and details are in Supplementary Methods.

### Methods to enhance detection of pathogen nucleic acid

We first assessed whether enrichment for non-methylated microbial DNA would improve mNGS yield. We used samples from 12 subjects: 10 with clinically diagnosed DNA virus infections and 2 with clinically diagnosed Lyme disease. Samples underwent methylated DNA depletion (MDD) using the NEBNext Microbiome DNA Enrichment Kit (New England Biolabs), and the enriched fraction was used for DNA library construction as above (Figure 1). Next, to assess the efficacy of enrichment for viral nucleic acid, we performed hybrid capture (HC) using a set of probes targeting all viruses known to infect humans (28). We applied HC to 13 RNA and 12 DNA libraries from subjects with clinically diagnosed RNA and DNA virus infections, respectively. Given the observed efficacy of HC, we also applied HC to samples from 20 subjects in the “Unknown” group (using the RNA library, DNA library, or both depending on clinical suspicion for a specific pathogen). To perform HC, indexed libraries were pooled in groups of approximately 5 samples per reaction, then underwent hybridization and capture using the SeqCap EZ Enrichment Kit (Roche) with modifications described previously (28). HC libraries were pooled and sequenced as described above. Finally, we applied both MDD and HC to a subset of 12 samples from patients with known or suspected DNA virus infection.

### Metagenomic analysis pipeline

Illumina sequencing reads were demultiplexed via viral-ngs, quality filtered and read trimmed using Trimmomatic (29), and depleted of human reads via a comprehensive KrakenUniq (30) database. Resulting reads were de-duplicated and assembled into metagenomic contigs via metaSPAdes (31). Contigs were classified using a cascading BLAST scheme in which unclassified contigs at each stage passed to the next level of more intensive BLAST searches from MegaBLAST, BLASTn, to BLASTx (32, 33). Contigs and associated hits derived from water and negative control samples were aggregated into a contaminant database and used to further deplete the human-depleted reads (Figure 1C, Supplementary Figure 1).

Finally, the human- and contaminant-depleted reads were classified by KrakenUniq using the same comprehensive database as above. Reads classified as potentially human-pathogenic viruses were validated via BLAST, discarding any reads that were not concordantly classified by both methods. The counts of reads per taxa were normalized to sequencing depth as reads per million (RPM). Kaiju was run on depleted reads to explore divergent taxa hits, while viral-ngs was used to assemble genomes for a subset of viruses.

### Statistical analyses

Analyses were performed using Student’s t-test and the Mann-Whitney *U* test for normally and non-normally distributed continuous variables, respectively, and using the *χ*^2^ test for categorical variables.

### Funding source

The funders of the study had no role in study design, data collection, data analysis, data interpretation, or writing of the report.

#### Data Availability

Reads after QC filtering, trimming, and depletion of human reads via KrakenUniq to a comprehensive database including the human genome (GRChg38/hg38) and all human sequences from the BLAST NT database, are available in the NCBI Sequence Read Archive (SRA) under accession PRJNA668392. Supplementary tables are available at DOI: 10.6084/m9.figshare.13266506, and supplementary figures at DOI: 10.6084/m9.figshare.13266488.

## Results

### Clinical characteristics

Of the 68 adults enrolled, 63% (43/68) were male, subjects ranged in age from 24 to 86 years (median = 58 years (interquartile range (IQR) [39, 72], Table 1) and 25 (37%) were immunocompromised (Supplementary Methods). New England was the primary residence for all except one subject who lived in Florida. Altered mental status was described in 56% (38/68), while a minority had photophobia (24% (16/68)) or neck stiffness (26% (18/68)). Twenty subjects out of 68 (29%) were admitted to the intensive care unit, and in-hospital mortality was 6% (4/68).

**Table 1:**
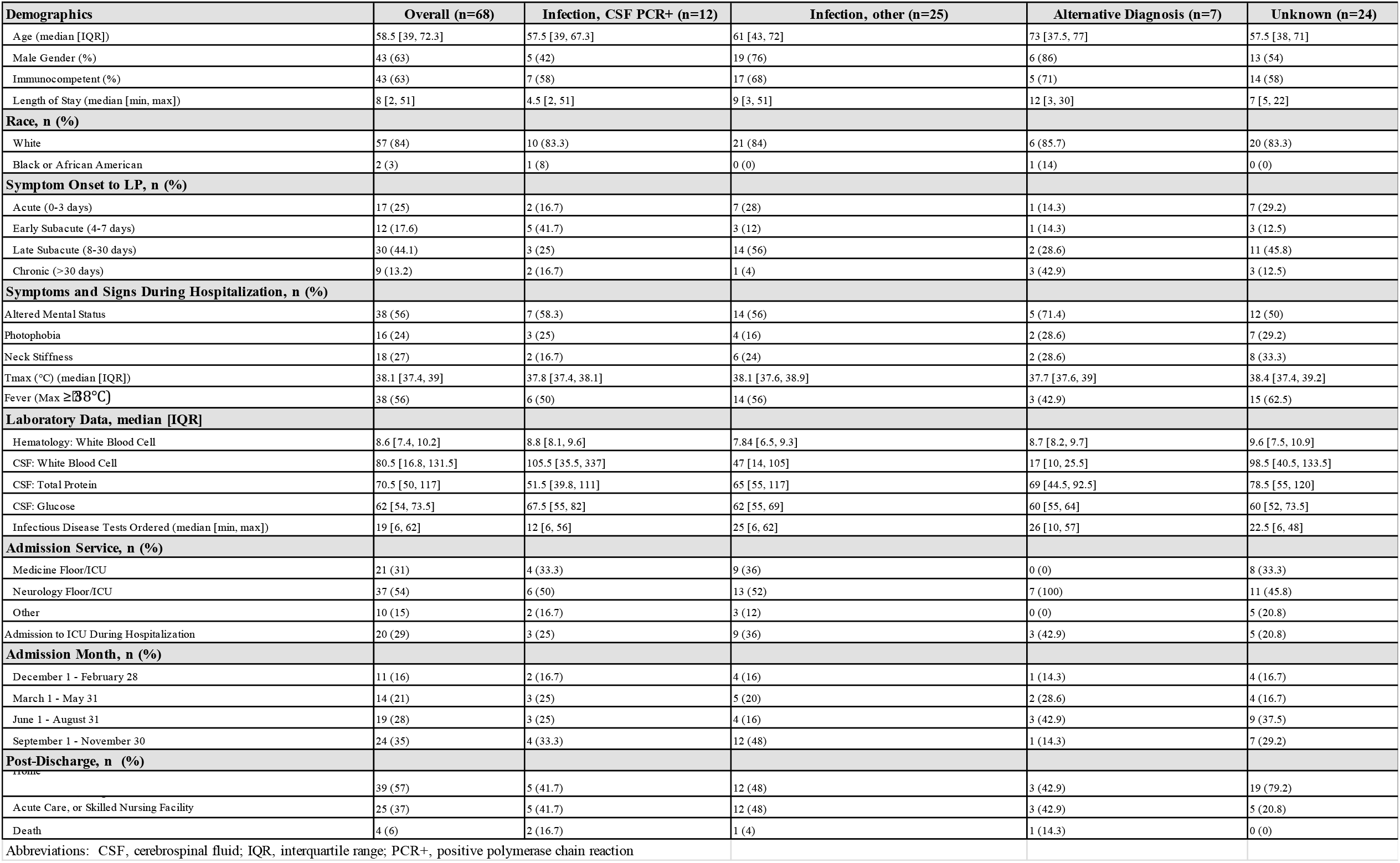
Clinical Characteristics of Enrolled Subjects Stratified by Diagnostic Groups.

Based on clinical testing, 44 of the 68 subjects received a conclusive diagnosis by discharge. Twelve subjects were diagnosed with viral infection by PCR from CSF (“Infection, CSF PCR+” group), 25 were diagnosed with infection by serology or PCR from blood (“Infection, Other” group), and 7 had a non-infectious etiology (“Alternative Diagnosis” group). The remaining 24 subjects (35%) had no known diagnosis (“Unknown” group) (Table 1). Subjects classified as “Unknown” underwent exhaustive clinical testing; 50% of them (12/24) had ≥ 25 infectious disease (ID) tests (Figure 2A; Supplementary Figure 2, Supplementary Table 2 (DOI: 10.6084/m9.figshare.13266506)), and no other diagnoses were made by clinical work-up alone in long-term follow-up (Supplementary Table 3, DOI: 10.6084/m9.figshare.13266506). In contrast, the “Infection, CSF PCR+” group had a much lower median number of clinical ID tests performed (12 [IQR:6,56)] vs (22·5 [IQR 11, 36)] for the “Unknown” group). The “Infection, CSF PCR+” group also had the shortest length of stay (LOS) (4·5 days [2,51)], and across the total cohort, LOS moderately correlated with the number of ID tests ordered (Spearman’s *ρ* = 0·65, p<0·01; Figure 2B).

**Figure 2.**
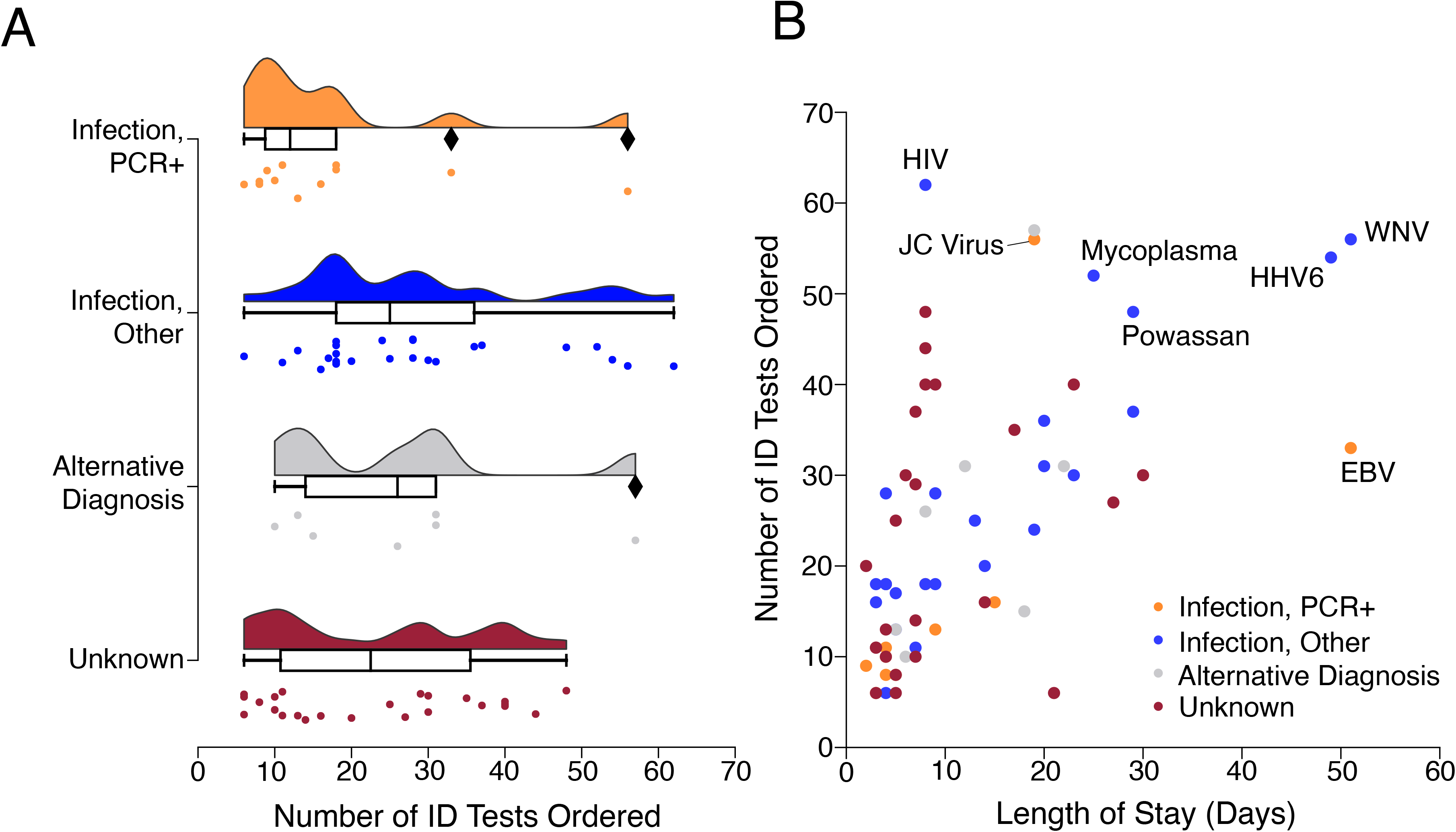
Number of infectious disease tests ordered and lengths of stay among subjects. Distributions showing the number of infectious disease (ID) related tests ordered per subject, stratified by clinical diagnosis category. ID tests were counted if ordered between hospital admission day one and hospital discharge. Box plots with horizontal bars represent medians and interquartile ranges for ID tests. Diamonds represent data points greater than 1.5 IQR (A). Scatter plot showing the number of ID tests versus length of stay per subject. Colors indicate clinical diagnosis category. LOS correlated with the number of total ID tests ordered (Spearman’s rho = 0·65, p<0·01). The final clinical diagnosis for viral pathogens is stated for cases whose number of ID tests or LOS was an outlier above the 3rd quartile (B).

### Results from mNGS and enhanced methods

To understand mNGS performance in a real-world context, we sequenced 68 CSF, 3 plasma, 5 serum, and 12 urine samples, along with 47 negative controls. We performed mNGS from RNA, DNA, or both, generating an average of 9.6 million reads per subject (Supplementary Figure 3, Supplementary Tables 4-6 (DOI: 10.6084/m9.figshare.13266506)). We identified a plausible pathogen in 22 subjects (32.4%): 18 by standard mNGS, an additional 2 with the use of HC, and 2 more with the use of MDD (Figure 3, Supplementary Figure 4). As expected, we detected viral nucleic acid in most subjects in the “Infection, CSF PCR+” group (10 out of 12, 83%, Figure 3), consistent with other mNGS studies (18, 23). mNGS was negative in one subject with herpes simplex virus 2 (HSV-2) and another with human immunodeficiency virus 1 (HIV-1), illustrating that mNGS can be less sensitive than PCR for very low-level infections (Supplementary Results, Supplementary Figure 5). We detected reads from both JC virus and HIV in a subject with HIV and progressive multifocal leukoencephalopathy (PML), illustrating the capacity of this single platform to identify viral coinfections. In assessing our enhanced methods, we found that HC increased the number of viral reads in 8 out of 9 cases positive by routine mNGS, sometimes substantially (Figure 4). By contrast, MDD led to mixed results, enabling virus detection in some cases (e.g., varicella zoster virus (VZV) in M049 and M070) while decreasing yield in others (e.g., Epstein-Barr virus (EBV) in M095) (Figure 4, Supplementary Results).

**Figure 3.**
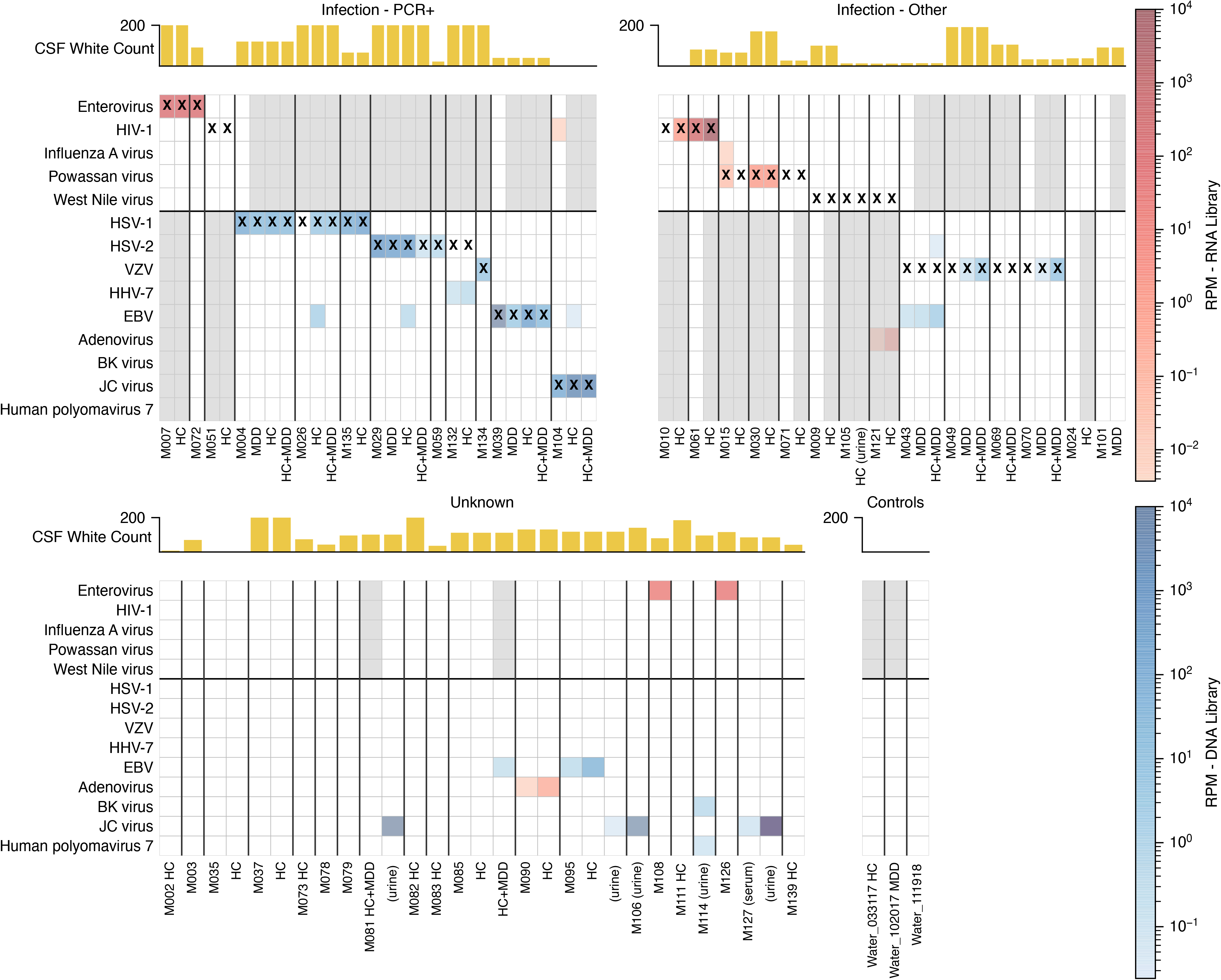
Viral taxa identified in cerebrospinal fluid using mNG with or without enhanced methods. Heatmap shows viral taxa identified in each sample. Rows are viral taxa, and columns are samples, some with enhanced sequencing methods (HC and/or MDD). Only classifications with over 100 unique kmers, at least 1 BLAST confirmed read, and manually reviewed as non-contaminant are shown. Rows are grouped by RNA viruses (top section) or DNA viruses (bottom section). Color intensity corresponds to the RPM of the taxa. Red boxes correspond to detection in RNA libraries while blue boxes correspond to detection in DNA libraries. Some DNA viruses were detected in RNA libraries (e.g. adenovirus in M121). Gray shaded columns represent samples that did not undergo DNA or RNA sequencing. Samples in which a contaminant were found are included here as blank columns, and the contaminants are shown in Supplementary Figure 4. Stars represent the clinical diagnosis. Yellow bars indicate the CSF nucleated cell count for each subject. The four groupings of columns from top left to bottom right correspond to infections diagnosed with a positive PCR, infections diagnosed by non-molecular techniques, subjects with unknown etiology, and negative controls including extracted water.

**Figure 4.**
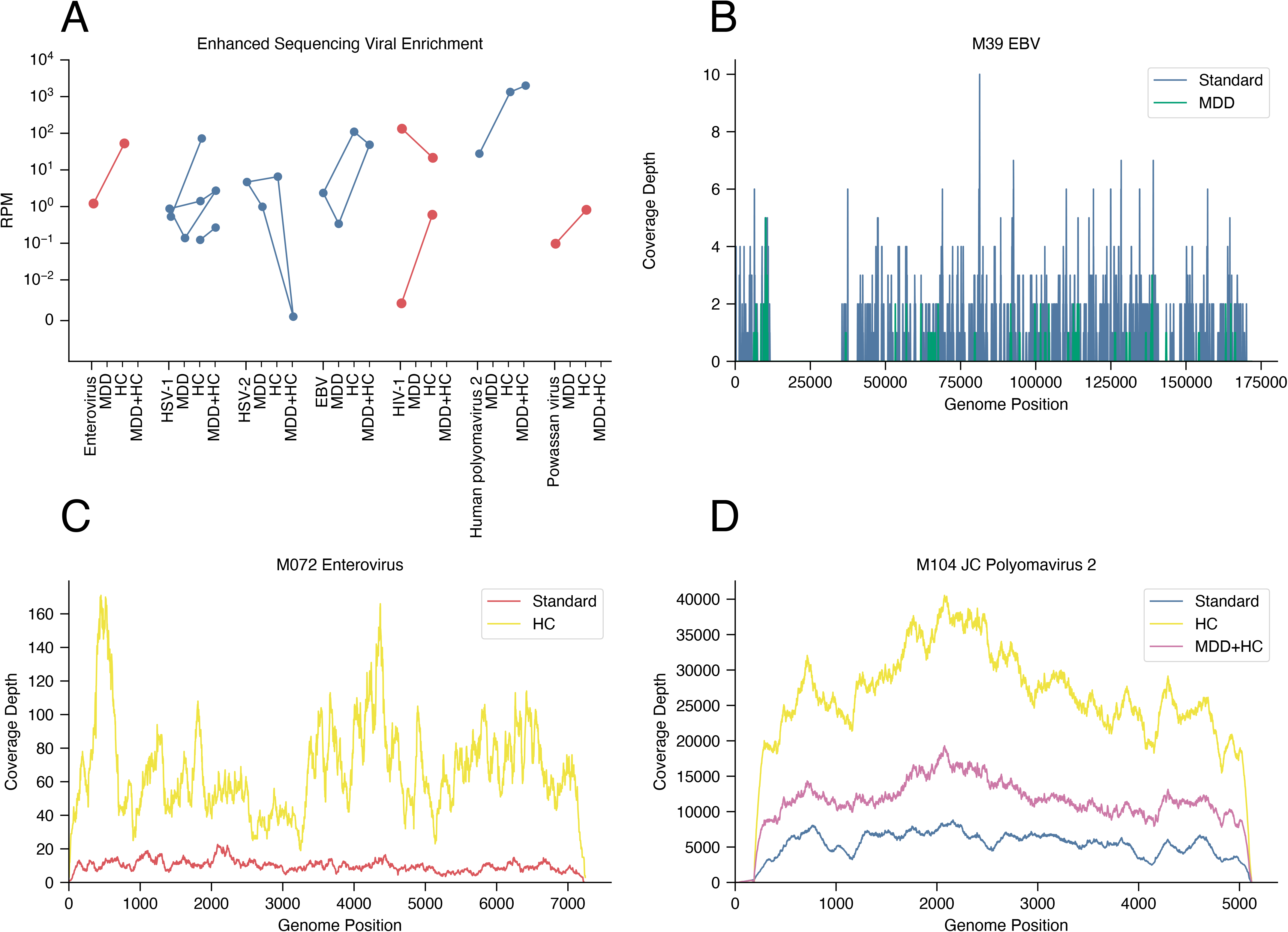
Enhanced methods for Metagenomic Next Generation Sequencing. Comparison of specific viral abundances among the non-computationally depleted reports for HC, MDD, and HC+MDD for RNA samples (orange) and DNA samples (blue) (A). Hybrid capture improved overall coverage for DNA and RNA viruses such as EBV (B), Enterovirus (C), and JC polyomavirus 2 (D). Methylated DNA depletion improved coverage for some DNA viruses such as JC polyomavirus 2 (panel D) but not others such as EBV, which utilizes host methylation in its life cycle (B).

### mNGS detects pathogens not traditionally diagnosed by CSF PCR

Twenty-five subjects in the “Infection, Other” group had infections diagnosed by serology from CSF and/or blood, or PCR from blood (Table 1). Fifteen had an infection for which no clinically approved CSF PCR assay was available; standard mNGS detected pathogen nucleic acid in six, and mNGS plus HC in a seventh, yielding 7/15 positive hits (47%) (Figure 3). There were several cases of regional interest. For example, three subjects were clinically diagnosed with Powassan encephalitis using a time-consuming send-out serology test, and mNGS identified Powassan virus RNA in two cases. In addition, while our methods were focused on viral detection, we identified atypical bacteria whose genome reads were readily distinguishable from background, including *Borrelia burgdorferi* in two out of two subjects diagnosed with Lyme disease by serology, and *Anaplasma phagocytophilum* in a subject diagnosed by PCR from blood (Supplementary Figure 6).

In the remaining ten subjects from the “Infection, Other” group, a clinical CSF PCR test was available and negative for the culprit pathogen (human herpesvirus 6 (HHV-6) (n=1), VZV (n=3), West Nile virus (WNV) (n=3), and mycoplasma (n=3)). While these were negative using standard mNGS, the addition of MDD allowed detection of VZV in two subjects (M049, M070) (Supplementary Results; Supplementary Figure 7). In both cases, clinical CSF VZV PCR from the same sample was negative, illustrating that mNGS may occasionally be more sensitive than a clinically validated PCR. By contrast, MDD decreased yield for other herpesviruses, suggesting pathogen-specific effects (Supplementary Results; Figure 4).

### mNGS detects pathogens not tested by clinicians

Among the 24 subjects with no identified clinical diagnosis (“Unknown”), standard mNGS identified viruses in three subjects, and no additional pathogens were detected using MDD and HC. We detected enterovirus in two subjects with lymphocytic meningitis (M108, M126), neither of whom had clinical enterovirus PCR testing. These findings were verified by sequencing a second CSF aliquot and by assembling a complete enterovirus genome for each subject. Phylogenetic analysis from both subjects demonstrated closely-related echovirus 30 strains (Supplementary Figure 8).

We also detected EBV and assembled a complete genome in one subject (M095) during two serial hospitalizations for recurrent lymphocytic meningitis. While clinical testing for EBV in CSF was not performed, EBV PCR was positive from blood during both admissions. Overall, these results are compatible with EBV meningitis or reactivation in the setting of another, unidentified primary syndrome (34).

### mNGS detects viruses of uncertain significance

In addition to the plausible pathogens described above, we detected DNA viruses of uncertain clinical significance. EBV was present at low levels in CSF from four subjects, three of whom had alternative primary diagnoses: VZV (M043), herpes simplex virus 1 (HSV-1) (M026), and HSV-2 (M029). For the fourth subject (M085), no alternative diagnosis was identified; however, EBV reads were only detected after MDD and HC, and a clinical PCR test for EBV from CSF was negative. Review of clinical data for these subjects suggested that EBV was unlikely to explain their clinical syndromes, and these findings most likely suggest reactivation in the setting of another acute process.

We also detected human herpesvirus 7 (HHV-7) at a low level in a subject (M132) who was diagnosed with HSV-2 by clinical PCR, but HSV-2 was not detected by mNGS. Acute encephalitis due to HHV-7 rarely occurs in immunocompetent adults, and has been described in three cases of patients with limbic encephalitis (35), facial cranial palsy, and polymyeloradiculitis (36, 37); none of these syndromes were compatible with this subject’s presentation. Adenovirus reads were detected in two subjects (M090, M121) and were not considered vector contaminants due to their distribution across the genome; however, the reads were found in RNA libraries only, and subjects were not known to be immunocompromised or have features compatible with adenovirus infection.

A known challenge of mNGS is assessment and interpretation of background contamination. Even after extensive computational depletion of both human reads and sequences found in negative controls, bacteria accounted for ~11% of DNA and ~39% of RNA reads. We also found viral reads from bacteriophages and vectors commonly used in molecular biology, such as adenovirus, cytomegalovirus (CMV), HIV/lentiviruses, and parvoviruses, consistent with prior studies (38). Finally, we found a handful of reads matching recently discovered picornaviruses from environmental surveys (Supplementary Results and Supplementary Table 7 (DOI: 10.6084/m9.figshare.13266506)) (39).

### mNGS is negative in subjects with non-infectious diagnosis

mNGS did not detect pathogen nucleic acid in the seven subjects with non-infectious diseases (“Alternative Diagnosis” group): autoimmune encephalitis and cerebellitis (n=3), lymphoma (n=2), and vasculitis (n=2). In this category, the median CSF WBC was 2-6 times lower than in the two infection groups. The “Alternative Diagnosis” group had the highest number of ID tests ordered in CSF and blood (median 26 tests, range [10,57]; Supplementary Table 2 (DOI: 10.6084/m9.figshare.13266506)) and is consistent with provider practice to test a wide-range of pathogens prior to immunomodulatory therapy; subjects were ultimately treated with immunosuppressive agents.

## Discussion

Advances in genomic technologies provide translational researchers the unprecedented capacity to identify and study pathogens in patients with meningitis and encephalitis. Here, we performed a prospective study using mNGS, enhanced laboratory and analysis techniques, and detailed clinical phenotyping to assess the use of this technology as a diagnostic tool for hospitalized subjects with inflammatory CSF. We identified a range of CNS pathogens, including regionally important tick-borne organisms not typically diagnosed by CSF nucleic acid testing. In 9 cases, we were able to recover full or partial viral genomes, demonstrating the utility of this technique for virus characterization studies (e.g., molecular epidemiology and identification of neurotropic variants). We also demonstrated that subjects with CNS infections diagnosed using CSF PCR assays undergo fewer ID tests compared to other clinical groups with inflammatory CSF and typically have shorter lengths of hospital stay (40); from this, we infer that the expansion of molecular diagnostic techniques such as mNGS may have direct and positive impacts on patient care and associated costs. Together with recent reports (18), this work highlights that mNGS is likely to become a mainstay in the infectious disease diagnostic toolkit.

Overall, mNGS was highly effective at detecting pathogens diagnosed by clinical PCR testing. Metagenomic NGS detected the expected pathogen in 10 of 12 subjects, similar to a recent study detecting viruses in 14 out of 16 subjects diagnosed by CSF PCR (18). Our results also highlight the benefit of enhanced mNGS techniques. For example, MDD plus mNGS detected VZV DNA in two additional samples negative by standard mNGS. However, MDD decreased yield for some viruses, indicating that the role of this technique in mNGS remains unclear. Pan-viral HC consistently improved sequencing of RNA and DNA viruses, and resulted in virus detection in two cases (HSV-1 and HIV) that were negative by standard mNGS.

An additional strength of this study was the detection of pathogens not routinely diagnosed by CSF PCR, most notably the tick-borne pathogens Powassan virus (17), *Borrelia burgdorferi,* and *Anaplasma phagocytophilum*. These pathogens have increasing rates of human infection (41), particularly in the Northeast U.S., where this study was conducted. For Powassan virus, which is routinely diagnosed by serology, our findings illustrate the potential utility of nucleic-acid based screening. Interestingly, we detected the CSF presence of *Anaplasma,* which is not commonly considered to be a cause of CNS infection (42), although the related intracellular bacteria *Ehrlichia chaffeensis* can cause meningoencephalitis (43, 44). Overall, the high number of subjects with tick-borne infection highlights the importance of conducting mNGS in diverse geographical regions for both diagnostic purposes and epidemiological studies.

Among the 24 subjects in whom no diagnosis was achieved by routine clinical testing (“Unknown”), metagenomic sequencing detected potential pathogens in three (8%), a rate similar to that of Wilson *et al* (13/159 = 8%) (18). It is possible that subjects in whom no pathogen nucleic acid was detected had a non-infectious syndrome or an infection with a low pathogen burden or short duration of replication. We reviewed the post-discharge clinical course in the subgroup and none were identified as having an infectious syndrome, signalling the likelihood that mNGS did not miss an actionable result.

Our results highlight a few challenges associated with mNGS, particularly for infections with low titer or para-infectious complications. For example, we report an equivocal mNGS result in a subject with HIV-1 who had a CSF HIV viral load of 469 copies/ml, a value close to the recently reported CSF limit of detection of 313 copies/ml for HIV-1 using mNGS (23). Additionally, mNGS results were negative in all four subjects with WNV, three of whom had clinical WNV PCR tests from CSF performed, which were also negative. These results support other studies showing that WNV nucleic acid is usually undetectable in CSF by clinical PCR (23, 45) or mNGS (18), though it may be observed in immunocompromised subjects (15, 45–48). Similarly, CSF mycoplasma nucleic acid was not detected clinically or by mNGS from three subjects with positive mycoplasma serology, supporting interpretations that CNS complications of mycoplasma infections likely reflect a para-infectious antibody-mediated response, rather than direct infection (49).

While we investigated specific atypical bacteria of interest (*Borrelia, Anaplasma, Mycoplasma* spp.), our study focused on viruses because they are the most common pathogen in CNS infection (4, 5, 24, 25), bacteria and fungi often require different laboratory methods for processing and nucleic acid extraction (50), bacterial infections are associated with greater pleocytosis and therefore higher levels of host background (23), and the analysis of viruses is more tractable given that mNGS (50) commonly detects bacterial reads (e.g., *P. aeruginosa, E. coli*) as background from skin and reagents (50). As this was not a clinical validation study, we focused on the practical application of mNGS in a defined cohort, rather than general diagnostic test performance (21, 23). We adhered to strict practices to minimize contamination but we did not conduct this research study in a Clinical Laboratory Improvement Amendments (CLIA)-certified laboratory (22), allowing us flexibility in iterative testing and refinement of methods. Because this study was conducted primarily using clinical excess samples, many of which had undergone multiple prior freeze-thaw cycles for clinical testing, it is also possible that some infections were missed due to nucleic acid degradation prior to mNGS, which would be solved if clinical processing for mNGS is standardized (50).

### Conclusions

Overall, our results highlight several important benefits of mNGS, including faster turn-around times than serology (17), recovery of pathogen genomic data, and reducing the dependence on test-specific diagnostics. However, our results among subjects with unknown etiology of disease suggest that the addition of mNGS to standard clinical testing will lead to relatively few additional diagnoses, underscoring the challenge of identifying an etiology in these devastating clinical syndromes. One potential strategy for incorporating mNGS into clinical diagnostic workflows would be wide implementation early in the diagnostic workup to capitalize on one-step detection of common pathogens, potentially sparing subjects unnecessary tests and reducing overall cost. An alternative would be to reserve this specialized technique for subjects with a high pre-test probability of infection (e.g., immunocompromised). Determining how to best utilize mNGS in clinical practice will require evaluation of these factors as well as the cost and logistics of implementation (51, 52). Currently, it is prudent to employ diagnostic mNGS through close communication between clinicians and mNGS experts (18) to evaluate the plausibility of pathogens identified. This is especially important considering background reads and contamination, the essential limitation that mNGS only detects infections with circulating pathogen nucleic acid, and our still-evolving understanding of mNGS test characteristics. Results from this study will inform ongoing efforts to transition the much-needed and promising technique of mNGS from a research tool to a clinical test used in the routine care of patients with suspected CNS infection.

## Acknowledgements

We are grateful to the patients, their families, and all medical staff involved in their care. We thank Graham McGrath for laboratory support, and Janice Stefanski and the Microbiology staff at MGH for facilitating sample acquisition.

## Author Contributions

Anne Piantadosi - study design, IRB site principal investigator, data collection, figures, data analyses, data interpretation, writing

Shibani Mukerji - study design, IRB principal investigator, data collection, figures, data analyses, data interpretation, writing

Simon Ye - figures, computational analyses, data interpretation, writing

Michael Leone - data collection, figures, literature search

Lisa Freimark - data collection

Daniel Park - data interpretation, data analyses

Gordon Adams - data collection

Jacob Lemieux - study design, data collection, data interpretation

Sanjat Kanjilal - data collection, data interpretation

Isaac Solomon - figures, data interpretation, writing

Asim Ahmed - data interpretation, writing

Robert Goldstein - study design, data analysis

Vijay Ganesh - data collection

Bridget Ostrem - data collection, data interpretation

Kaeyln Cummins - figures

Jesse Thon - writing

Cormac Kinsella - data analysis, data interpretation

Eric Rosenberg - study design, data interpretation

Matthew Frosch - study design, data interpretation

Marcia Goldberg - study design, data interpretation

Tracey Cho - study design, data interpretation, writing

Pardis Sabeti - study design, data interpretation, writing

---

**Supplementary Figure 1. Computational processing workflow**

Sequencing reads first underwent universal quality control, human depletion (via stringent criteria of >20% kmers within the read classifying specifically to human taxid), and de-duplication (A). These reads were assembled into contigs, and >600bp contigs were BLASTed to recover strong reference matches for long contigs (B). These were used as a “negative controls” depletion database, after which remaining reads were classified via comprehensive Krakenuniq and Kaiju databases. Viral hits were validated using BLASTn.

**Supplementary Figure 2. Correlations between length of hospitalization and diagnostic testing ordered, stratified by clinical diagnosis.**

Box plots show median length of stay (LOS; horizontal line) and whiskers indicate 1st and 3rd quartile. Dots indicate a LOS greater than 1.5*Interquartile range. There were no significant differences in LOS between clinical diagnosis groups (A). Scatter plot showing the number of CSF tests versus LOS (B) and number of PCR tests versus LOS (C). Colors indicate clinical diagnosis category. LOS moderately correlated with the number of total ID tests ordered (Spearman’s *ρ* = 0·65, p<0·01; Figure 2B), with number of tests ordered from CSF only (Spearman’s *ρ* = 0·46, p<0·01; Supplementary Figure 2B)

**Supplementary Figure 3. Sequencing metrics for various stages of the computational pipeline**

The total number of reads in each sequencing library from raw de-multiplexed reads through the stages of quality control/trimming, human depletion, deduplication, and negative depletion (A). The distribution of the percentage of reads retained after each incremental step for all samples (C). Comparison of human abundance for each subject between routine, hybrid capture (HC), methylated DNA depletion (MDD), and hybrid capture plus methylated DNA depletion (HC+MDD) on DNA samples (B). Comparison for RNA (D).

**Supplementary Figure 4: Unfiltered metagenomic classifications including contaminants**

Heatmap shows viral taxa identified in each sample type. Compared to Figure 3, this Figure shows all classified taxa without manually screening out contaminants. Rows are viral taxa, and columns are sample types, some with enhanced sequencing methods (HC and/or MDD). Only classifications with over 100 unique kmers, and at least 1 BLAST confirmed read are shown. Rows are grouped by whether they are RNA viruses vs DNA viruses (top vs bottom section). Color intensity corresponds to the RPM of the taxa. Red boxes correspond to detection in RNA libraries while blue boxes correspond to detection in DNA libraries. Stars represent the clinical diagnosis. Gray shaded columns represent samples that did not undergo DNA or RNA sequencing. The yellow bars indicate nucleated cell count in the CSF for each subject. The four groupings of columns from top left to bottom right correspond to infections diagnosed with a positive PCR, infections diagnosed by non-molecular techniques, subjects with unknown etiology, and negative controls including water.

**Supplementary Figure 5. Results of HSV-2 and HIV-1 specific PCR**

Amplification curve analysis demonstrated that CSF from subject M029 (blue, positive control) amplified in three out of three replicates (mean Ct = 23.8), consistent with positive mNGS results for HSV-2. CSF from subject M132 (gray) amplified in only one out of three replicates (Ct = 39.8); correspondingly, no HSV-2 reads were detected by mNGS. There was no amplification from the negative control (red) (A). Melting curve analysis demonstrated consistent curves across all positive wells(B). Amplification curve analysis demonstrated that CSF from subject M061 (purple, positive control) amplified in three out of three replicates (mean Ct = 25.2), consistent with positive mNGS results for HIV-1. CSF from subjects M051 (blue) and M010 (gray) amplified at high Ct values, similar to the negative control (red) (C). Melting curve analysis demonstrated that only one replicate from M051 melted in a pattern consistent with the positive controls; the other positive wells melted at lower temperatures, suggestive of nonspecific amplification or primer-dimerization (D). Gel electrophoresis results from PCR products demonstrate a band of the expected size for subject M061 (positive control) and a faint band of the expected size for subject M051, but not M010 or the negative control (E).

**Supplementary Figure 6. Enterovirus phylogeny**

Enterovirus genomes assembled from subjects in this study (red) were aligned with representative reference sequences for each subtype within the enterovirus B species (blue). This allowed classification of viral subtypes as Coxsackie B4 for M007 and echovirus 30 for M072, M108, and M126. Interestingly, viruses from M108 and M126, who were admitted approximately one month apart from one another and had no known epidemiological links, differed by only 0.6% (42 nucleotides), suggesting a common local circulating strain. Abbreviations: EV = enterovirus; Echo = echovirus; Cox = coxsackievirus.

**Supplementary Figure 7. Clinical course, laboratory findings and mNGS for subjects diagnosed with Varicella Zoster Virus**

Eight subjects were diagnosed clinically with varicella zoster virus (VZV)-related neurological diseases. Cerebrospinal fluid (CSF) metagenomic next generation sequencing (mNGS) was positive in three cases of VZV meningoencephalitis (red bar), and negative for the other five subjects. Positive cases had acute onset of symptoms and no prior antiviral treatment or minimal exposure (treatment and symptoms bars). CSF white blood cells at time of clinical VZV testing and mNGS VZV results are shown.

**Supplementary Figure 8: Detection of atypical bacteria**

Heatmap shows the recovery of sequencing reads for a subset of atypical bacteria. Only samples classified with over 100 unique kmers, and at least 1 BLAST confirmed read are shown. Color intensity corresponds to RPM in DNA samples.

## References

1. Feigin VL, Nichols E, Alam T, Bannick MS, Beghi E, Blake N, Culpepper WJ, Dorsey ER, Elbaz A, Ellenbogen RG, Fisher JL, Fitzmaurice C, Giussani G, Glennie L, James SL, Johnson CO, Kassebaum NJ, Logroscino G, Marin B, Mountjoy-Venning WC, Nguyen M, Ofori-Asenso R, Patel AP, Piccininni M, Roth GA, Steiner TJ, Stovner LJ, Szoeke CEI, Theadom A, Vollset SE, Wallin MT, Wright C, Zunt JR, Abbasi N, Abd-Allah F, Abdelalim A, Abdollahpour I, Aboyans V, Abraha HN, Acharya D, Adamu AA, Adebayo OM, Adeoye AM, Adsuar JC, Afarideh M, Agrawal S, Ahmadi A, Ahmed MB, Aichour AN, Aichour I, Aichour MTE, Akinyemi RO, Akseer N, Al-Eyadhy A, Al-Shahi Salman R, Alahdab F, Alene KA, Aljunid SM, Altirkawi K, Alvis-Guzman N, Anber NH, Antonio CAT, Arabloo J, Aremu O, Ärnlöv J, Asayesh H, Asghar RJ, Atalay HT, Awasthi A, Ayala Quintanilla BP, Ayuk TB, Badawi A, Banach M, Banoub JAM, Barboza MA, Barker-Collo SL, Bärnighausen TW, Baune BT, Bedi N, Behzadifar M, Behzadifar M, Béjot Y, Bekele BB, Belachew AB, Bennett DA, Bensenor IM, Berhane A, Beuran M, Bhattacharyya K, Bhutta ZA, Biadgo B, Bijani A, Bililign N, Bin Sayeed MS, Blazes CK, Brayne C, Butt ZA, Campos-Nonato IR, Cantu-Brito C, Car M, Cárdenas R, Carrero JJ, Carvalho F, Castañeda-Orjuela CA, Castro F, Catalá-López F, Cerin E, Chaiah Y, Chang J-C, Chatziralli I, Chiang PP-C, Christensen H, Christopher DJ, Cooper C, Cortesi PA, Costa VM, Criqui MH, Crowe CS, Damasceno AAM, Daryani A, De la Cruz-Góngora V, De la Hoz FP, De Leo D, Demoz GT, Deribe K, Dharmaratne SD, Diaz D, Dinberu MT, Djalalinia S, Doku DT, Dubey M, Dubljanin E, Duken EE, Edvardsson D, El-Khatib Z, Endres M, Endries AY, Eskandarieh S, Esteghamati A, Esteghamati S, Farhadi F, Faro A, Farzadfar F, Farzaei MH, Fatima B, Fereshtehnejad S-M, Fernandes E, Feyissa GT, Filip I, Fischer F, Fukumoto T, Ganji M, Gankpe FG, Garcia-Gordillo MA, Gebre AK, Gebremichael TG, Gelaw BK, Geleijnse JM, Geremew D, Gezae KE, Ghasemi-Kasman M, Gidey MY, Gill PS, Gill TK, Girma ET, Gnedovskaya EV, Goulart AC, Grada A, Grosso G, Guo Y, Gupta R, Gupta R, Haagsma JA, Hagos TB, Haj-Mirzaian A, Haj-Mirzaian A, Hamadeh RR, Hamidi S, Hankey GJ, Hao Y, Haro JM, Hassankhani H, Hassen HY, Havmoeller R, Hay SI, Hegazy MI, Heidari B, Henok A, Heydarpour F, Hoang CL, Hole MK, Homaie Rad E, Hosseini SM, Hu G, Igumbor EU, Ilesanmi OS, Irvani SSN, Islam SMS, Jakovljevic M, Javanbakht M, Jha RP, Jobanputra YB, Jonas JB, Jozwiak JJ, Jürisson M, Kahsay A, Kalani R, Kalkonde Y, Kamil TA, Kanchan T, Karami M, Karch A, Karimi N, Kasaeian A, Kassa TD, Kassa ZY, Kaul A, Kefale AT, Keiyoro PN, Khader YS, Khafaie MA, Khalil IA, Khan EA, Khang Y-H, Khazaie H, Kiadaliri AA, Kiirithio DN, Kim AS, Kim D, Kim Y-E, Kim YJ, Kisa A, Kokubo Y, Koyanagi A, Krishnamurthi RV, Kuate Defo B, Kucuk Bicer B, Kumar M, Lacey B, Lafranconi A, Lansingh VC, Latifi A, Leshargie CT, Li S, Liao Y, Linn S, Lo WD, Lopez JCF, Lorkowski S, Lotufo PA, Lucas RM, Lunevicius R, Mackay MT, Mahotra NB, Majdan M, Majdzadeh R, Majeed A, Malekzadeh R, Malta DC, Manafi N, Mansournia MA, Mantovani LG, März W, Mashamba-Thompson TP, Massenburg BB, Mate KKV, McAlinden C, McGrath JJ, Mehta V, Meier T, Meles HG, Melese A, Memiah PTN, Memish ZA, Mendoza W, Mengistu DT, Mengistu G, Meretoja A, Meretoja TJ, Mestrovic T, Miazgowski B, Miazgowski T, Miller TR, Mini GK, Mirrakhimov EM, Moazen B, Mohajer B, Mohammad Gholi Mezerji N, Mohammadi M, Mohammadi-Khanaposhtani M, Mohammadibakhsh R, Mohammadnia-Afrouzi M, Mohammed S, Mohebi F, Mokdad AH, Monasta L, Mondello S, Moodley Y, Moosazadeh M, Moradi G, Moradi-Lakeh M, Moradinazar M, Moraga P, Moreno Velásquez I, Morrison SD, Mousavi SM, Muhammed OS, Muruet W, Musa KI, Mustafa G, Naderi M, Nagel G, Naheed A, Naik G, Najafi F, Nangia V, Negoi I, Negoi RI, Newton CRJ, Ngunjiri JW, Nguyen CT, Nguyen LH, Ningrum DNA, Nirayo YL, Nixon MR, Norrving B, Noubiap JJ, Nourollahpour Shiadeh M, Nyasulu PS, Ogah OS, Oh I-H, Olagunju AT, Olagunju TO, Olivares PR, Onwujekwe OE, Oren E, Owolabi MO, Pa M, Pakpour AH, Pan W-H, Panda-Jonas S, Pandian JD, Patel SK, Pereira DM, Petzold M, Pillay JD, Piradov MA, Polanczyk GV, Polinder S, Postma MJ, Poulton R, Poustchi H, Prakash S, Prakash V, Qorbani M, Radfar A, Rafay A, Rafiei A, Rahim F, Rahimi-Movaghar V, Rahman M, Rahman MHU, Rahman MA, Rajati F, Ram U, Ranta A, Rawaf DL, Rawaf S, Reinig N, Reis C, Renzaho AMN, Resnikoff S, Rezaeian S, Rezai MS, Rios González CM, Roberts NLS, Roever L, Ronfani L, Roro EM, Roshandel G, Rostami A, Sabbagh P, Sacco RL, Sachdev PS, Saddik B, Safari H, Safari-Faramani R, Safi S, Safiri S, Sagar R, Sahathevan R, Sahebkar A, Sahraian MA, Salamati P, Salehi Zahabi S, Salimi Y, Samy AM, Sanabria J, Santos IS, Santric Milicevic MM, Sarrafzadegan N, Sartorius B, Sarvi S, Sathian B, Satpathy M, Sawant AR, Sawhney M, Schneider IJC, Schöttker B, Schwebel DC, Seedat S, Sepanlou SG, Shabaninejad H, Shafieesabet A, Shaikh MA, Shakir RA, Shams-Beyranvand M, Shamsizadeh M, Sharif M, Sharif-Alhoseini M, She J, Sheikh A, Sheth KN, Shigematsu M, Shiri R, Shirkoohi R, Shiue I, Siabani S, Siddiqi TJ, Sigfusdottir ID, Sigurvinsdottir R, Silberberg DH, Silva JP, Silveira DGA, Singh JA, Sinha DN, Skiadaresi E, Smith M, Sobaih BH, Sobhani S, Soofi M, Soyiri IN, Sposato LA, Stein DJ, Stein MB, Stokes MA, Sufiyan MB, Sykes BL, Sylaja PN, Tabarés-Seisdedos R, Te Ao BJ, Tehrani-Banihashemi A, Temsah M-H, Temsah O, Thakur JS, Thrift AG, Topor-Madry R, Tortajada-Girbés M, Tovani-Palone MR, Tran BX, Tran KB, Truelsen TC, Tsadik AG, Tudor Car L, Ukwaja KN, Ullah I, Usman MS, Uthman OA, Valdez PR, Vasankari TJ, Vasanthan R, Veisani Y, Venketasubramanian N, Violante FS, Vlassov V, Vosoughi K, Vu GT, Vujcic IS, Wagnew FS, Waheed Y, Wang Y-P, Weiderpass E, Weiss J, Whiteford HA, Wijeratne T, Winkler AS, Wiysonge CS, Wolfe CDA, Xu G, Yadollahpour A, Yamada T, Yano Y, Yaseri M, Yatsuya H, Yimer EM, Yip P, Yisma E, Yonemoto N, Yousefifard M, Yu C, Zaidi Z, Zaman SB, Zamani M, Zandian H, Zare Z, Zhang Y, Zodpey S, Naghavi M, Murray CJL, Vos T. 2019. Global, regional, and national burden of neurological disorders, 1990–2016: a systematic analysis for the Global Burden of Disease Study 2016. Lancet Neurol 18:459–480.

2. McGill F, Griffiths MJ, Bonnett LJ, Geretti AM, Michael BD, Beeching NJ, McKee D, Scarlett P, Hart IJ, Mutton KJ, Jung A, Adan G, Gummery A, Sulaiman WAW, Ennis K, Martin AP, Haycox A, Miller A, Solomon T, UK Meningitis Study Investigators. 2018. Incidence, aetiology, and sequelae of viral meningitis in UK adults: a multicentre prospective observational cohort study. Lancet Infect Dis 18:992–1003.

3. Quist-Paulsen E, Ormaasen V, Kran A-MB, Dunlop O, Ueland PM, Ueland T, Eikeland R, Aukrust P, Nordenmark TH. 2019. Encephalitis and aseptic meningitis: short-term and long-term outcome, quality of life and neuropsychological functioning. Sci Rep 9:16158.

4. Glaser CA, Honarmand S, Anderson LJ, Schnurr DP, Forghani B, Cossen CK, Schuster FL, Christie LJ, Tureen JH. 2006. Beyond viruses: clinical profiles and etiologies associated with encephalitis. Clin Infect Dis 43:1565–1577.

5. Kupila L, Vuorinen T, Vainionpää R, Hukkanen V, Marttila RJ, Kotilainen P. 2006. Etiology of aseptic meningitis and encephalitis in an adult population. Neurology 66:75–80.

6. Nesher L, Hadi CM, Salazar L, Wootton SH, Garey KW, Lasco T, Luce AM, Hasbun R. 2016. Epidemiology of meningitis with a negative CSF Gram stain: under-utilization of available diagnostic tests. Epidemiol Infect 144:189–197.

7. Simner PJ, Miller S, Carroll KC. 2018. Understanding the Promises and Hurdles of Metagenomic Next-Generation Sequencing as a Diagnostic Tool for Infectious Diseases. Clin Infect Dis 66:778–788.

8. Gu W, Miller S, Chiu CY. 2019. Clinical Metagenomic Next-Generation Sequencing for Pathogen Detection. Annu Rev Pathol 14:319–338.

9. Chiu CY, Miller SA. 2019. Clinical metagenomics. Nat Rev Genet 20:341–355.

10. Wilson MR, Naccache SN, Samayoa E, Biagtan M, Bashir H, Yu G, Salamat SM, Somasekar S, Federman S, Miller S, Sokolic R, Garabedian E, Candotti F, Buckley RH, Reed KD, Meyer TL, Seroogy CM, Galloway R, Henderson SL, Gern JE, DeRisi JL, Chiu CY. 2014. Actionable diagnosis of neuroleptospirosis by next-generation sequencing. N Engl J Med 370:2408–2417.

11. Mongkolrattanothai K, Naccache SN. 2017. Neurobrucellosis: unexpected answer from metagenomic next-generation sequencing. Journal of the.

12. Fan S, Ren H, Wei Y, Mao C, Ma Z, Zhang L, Wang L, Ge Y, Li T, Cui L, Wu H, Guan H. 2018. Next-generation sequencing of the cerebrospinal fluid in the diagnosis of neurobrucellosis. Int J Infect Dis 67:20–24.

13. Christopeit M, Grundhoff A, Rohde H, Belmar-Campos C, Grzyska U, Fiehler J, Wolschke C, Ayuk F, Kröger N, Fischer N. 2016. Suspected encephalitis with Candida tropicalis and Fusarium detected by unbiased RNA sequencing. Ann Hematol 95:1919–1921.

14. Wilson MR, Shanbhag NM, Reid MJ, Singhal NS, Gelfand JM, Sample HA, Benkli B, O’Donovan BD, Ali IKM, Keating MK, Dunnebacke TH, Wood MD, Bollen A, DeRisi JL. 2015. Diagnosing Balamuthia mandrillaris Encephalitis With Metagenomic Deep Sequencing. Ann Neurol 78:722–730.

15. Wilson MR, Zimmermann LL, Crawford ED, Sample HA, Soni PR, Baker AN, Khan LM, DeRisi JL. 2017. Acute West Nile Virus Meningoencephalitis Diagnosed Via Metagenomic Deep Sequencing of Cerebrospinal Fluid in a Renal Transplant Patient. Am J Transplant 17:803–808.

16. Murkey JA, Chew KW, Carlson M, Shannon CL, Sirohi D, Sample HA, Wilson MR, Vespa P, Humphries RM, Miller S, Others. 2017. Hepatitis E Virus--associated meningoencephalitis in a lung transplant recipient diagnosed by clinical metagenomic sequencingOpen forum infectious diseases. Oxford University Press.

17. Piantadosi A, Kanjilal S, Ganesh V, Khanna A, Hyle EP, Rosand J, Bold T, Metsky HC, Lemieux J, Leone MJ, Freimark L, Matranga CB, Adams G, McGrath G, Zamirpour S, Telford S 3rd, Rosenberg E, Cho T, Frosch MP, Goldberg MB, Mukerji SS, Sabeti PC. 2018. Rapid Detection of Powassan Virus in a Patient With Encephalitis by Metagenomic Sequencing. Clin Infect Dis 66:789–792.

18. Wilson MR, Sample HA, Zorn KC, Arevalo S, Yu G, Neuhaus J, Federman S, Stryke D, Briggs B, Langelier C, Berger A, Douglas V, Josephson SA, Chow FC, Fulton BD, DeRisi JL, Gelfand JM, Naccache SN, Bender J, Dien Bard J, Murkey J, Carlson M, Vespa PM, Vijayan T, Allyn PR, Campeau S, Humphries RM, Klausner JD, Ganzon CD, Memar F, Ocampo NA, Zimmermann LL, Cohen SH, Polage CR, DeBiasi RL, Haller B, Dallas R, Maron G, Hayden R, Messacar K, Dominguez SR, Miller S, Chiu CY. 2019. Clinical Metagenomic Sequencing for Diagnosis of Meningitis and Encephalitis. N Engl J Med 380:2327–2340.

19. Haston JC, Rostad CA, Jerris RC, Milla SS, McCracken C, Pratt C, Wiley M, Prieto K, Palacios G, Shane AL, McElroy AK. 2019. Prospective Cohort Study of Next-Generation Sequencing as a Diagnostic Modality for Unexplained Encephalitis in Children. J Pediatric Infect Dis Soc https://doi.org/10.1093/jpids/piz032.

20. Brown JR, Bharucha T, Breuer J. 2018. Encephalitis diagnosis using metagenomics: application of next generation sequencing for undiagnosed cases. J Infect 76:225–240.

21. Schlaberg R, Chiu CY, Miller S, Procop GW, Weinstock G, Professional Practice Committee and Committee on Laboratory Practices of the American Society for Microbiology, Microbiology Resource Committee of the College of American Pathologists. 2017. Validation of Metagenomic Next-Generation Sequencing Tests for Universal Pathogen Detection. Arch Pathol Lab Med 141:776–786.

22. Blauwkamp TA, Thair S, Rosen MJ, Blair L, Lindner MS, Vilfan ID, Kawli T, Christians FC, Venkatasubrahmanyam S, Wall GD, Cheung A, Rogers ZN, Meshulam-Simon G, Huijse L, Balakrishnan S, Quinn JV, Hollemon D, Hong DK, Vaughn ML, Kertesz M, Bercovici S, Wilber JC, Yang S. 2019. Analytical and clinical validation of a microbial cell-free DNA sequencing test for infectious disease. Nature Microbiology.

23. Miller S, Naccache SN, Samayoa E, Messacar K, Arevalo S, Federman S, Stryke D, Pham E, Fung B, Bolosky WJ, Ingebrigtsen D, Lorizio W, Paff SM, Leake JA, Pesano R, DeBiasi R, Dominguez S, Chiu CY. 2019. Laboratory validation of a clinical metagenomic sequencing assay for pathogen detection in cerebrospinal fluid. Genome Res 29:831–842.

24. Glaser CA, Gilliam S, Schnurr D. 2003. In search of encephalitis etiologies: diagnostic challenges in the California Encephalitis Project, 1998—2000. Clin Infect Dis.

25. Vora NM, Holman RC, Mehal JM, Steiner CA, Blanton J, Sejvar J. 2014. Burden of encephalitis-associated hospitalizations in the United States, 1998–2010. Neurology 82:443–451.

26. Gire SK, Goba A, Andersen KG, Sealfon RSG, Park DJ, Kanneh L, Jalloh S, Momoh M, Fullah M, Dudas G, Wohl S, Moses LM, Yozwiak NL, Winnicki S, Matranga CB, Malboeuf CM, Qu J, Gladden AD, Schaffner SF, Yang X, Jiang P-P, Nekoui M, Colubri A, Coomber MR, Fonnie M, Moigboi A, Gbakie M, Kamara FK, Tucker V, Konuwa E, Saffa S, Sellu J, Jalloh AA, Kovoma A, Koninga J, Mustapha I, Kargbo K, Foday M, Yillah M, Kanneh F, Robert W, Massally JLB, Chapman SB, Bochicchio J, Murphy C, Nusbaum C, Young S, Birren BW, Grant DS, Scheiffelin JS, Lander ES, Happi C, Gevao SM, Gnirke A, Rambaut A, Garry RF, Khan SH, Sabeti PC. 2014. Genomic surveillance elucidates Ebola virus origin and transmission during the 2014 outbreak. Science 345:1369–1372.

27. Matranga CB, Andersen KG, Winnicki S, Busby M, Gladden AD, Tewhey R, Stremlau M, Berlin A, Gire SK, England E, Moses LM, Mikkelsen TS, Odia I, Ehiane PE, Folarin O, Goba A, Kahn SH, Grant DS, Honko A, Hensley L, Happi C, Garry RF, Malboeuf CM, Birren BW, Gnirke A, Levin JZ, Sabeti PC. 2014. Enhanced methods for unbiased deep sequencing of Lassa and Ebola RNA viruses from clinical and biological samples. Genome Biol 15:519.

28. Metsky HC, Siddle KJ, Gladden-Young A, Qu J, Yang DK, Brehio P, Goldfarb A, Piantadosi A, Wohl S, Carter A, Lin AE, Barnes KG, Tully DC, Corleis B, Hennigan S, Barbosa-Lima G, Vieira YR, Paul LM, Tan AL, Garcia KF, Parham LA, Odia I, Eromon P, Folarin OA, Goba A, Viral Hemorrhagic Fever Consortium, Simon-Lorière E, Hensley L, Balmaseda A, Harris E, Kwon DS, Allen TM, Runstadler JA, Smole S, Bozza FA, Souza TML, Isern S, Michael SF, Lorenzana I, Gehrke L, Bosch I, Ebel G, Grant DS, Happi CT, Park DJ, Gnirke A, Sabeti PC, Matranga CB. 2019. Capturing sequence diversity in metagenomes with comprehensive and scalable probe design. Nat Biotechnol 37:160–168.

29. Bolger AM, Lohse M, Usadel B. 2014. Trimmomatic: a flexible trimmer for Illumina sequence data. Bioinformatics 30:2114–2120.

30. Breitwieser FP, Baker DN, Salzberg SL. 2018. KrakenUniq: confident and fast metagenomics classification using unique k-mer counts. Genome Biol 19:198.

31. Nurk S, Meleshko D, Korobeynikov A, Pevzner PA. 2017. metaSPAdes: a new versatile metagenomic assembler. Genome Research.

32. Altschul SF, Gish W, Miller W, Myers EW, Lipman DJ. 1990. Basic local alignment search tool. J Mol Biol 215:403–410.

33. Morgulis A, Coulouris G, Raytselis Y, Madden TL, Agarwala R, Schäffer AA. 2008. Database indexing for production MegaBLAST searches. Bioinformatics 24:1757–1764.

34. Gilden DH, Mahalingam R, Cohrs RJ, Tyler KL. 2007. Herpesvirus infections of the nervous system. Nat Clin Pract Neurol 3:82–94.

35. Aburakawa Y, Katayama T, Saito T, Sawada J, Suzutani T, Aizawa H, Hasebe N. 2017. Limbic Encephalitis Associated with Human Herpesvirus-7 (HHV-7) in an Immunocompetent Adult: The First Reported Case in Japan. Intern Med 56:1919–1923.

36. Riva N, Franconi I, Meschiari M, Franceschini E, Puzzolante C, Cuomo G, Bianchi A, Cavalleri F, Genovese M, Mussini C. 2017. Acute human herpes virus 7 (HHV-7) encephalitis in an immunocompetent adult patient: a case report and review of literature. Infection 45:385–388.

37. Parra M, Alcala A, Amoros C, Baeza A, Galiana A, Tarragó D, García-Quesada MÁ, Sánchez-Hellín V. 2017. Encephalitis associated with human herpesvirus-7 infection in an immunocompetent adult. Virology Journal.

38. Asplund M, Kjartansdóttir KR, Mollerup S, Vinner L, Fridholm H, Herrera JAR, Friis-Nielsen J, Hansen TA, Jensen RH, Nielsen IB, Richter SR, Rey-Iglesia A, Matey-Hernandez ML, Alquezar-Planas DE, Olsen PVS, Sicheritz-Pontén T, Willerslev E, Lund O, Brunak S, Mourier T, Nielsen LP, Izarzugaza JMG, Hansen AJ. 2019. Contaminating viral sequences in high-throughput sequencing viromics: a linkage study of 700 sequencing libraries. Clinical Microbiology and Infection.

39. Shi M, Lin X-D, Tian J-H, Chen L-J, Chen X, Li C-X, Qin X-C, Li J, Cao J-P, Eden J-S, Buchmann J, Wang W, Xu J, Holmes EC, Zhang Y-Z. 2016. Redefining the invertebrate RNA virosphere. Nature 540:539–543.

40. Vareil M, Wille H, Kassab S, Le-Cornec C, Puges M, Desclaux A, Lafon ME, Tumiotto C, Cazanave C, Neau D. 2018. Clinical and biological features of enteroviral meningitis among adults and children and factors associated with severity and length of stay. J Clin Virol 104:56–60.

41. Rosenberg R, Lindsey NP, Fischer M, Gregory CJ, Hinckley AF, Mead PS, Paz-Bailey G, Waterman SH, Drexler NA, Kersh GJ, Hooks H, Partridge SK, Visser SN, Beard CB, Petersen LR. 2018. Vital Signs: Trends in Reported Vectorborne Disease Cases - United States and Territories, 2004-2016. MMWR Morb Mortal Wkly Rep 67:496–501.

42. Bennett MD J, Dolin R, Blaser MJ. 2014. Mandell, Douglas, and Bennett’s Principles and Practice of Infectious Diseases: 2-Volume Set. Elsevier Health Sciences.

43. Dahlgren FS, Mandel EJ, Krebs JW, Massung RF, McQuiston JH. 2011. Increasing incidence of Ehrlichia chaffeensis and Anaplasma phagocytophilum in the United States, 2000-2007. Am J Trop Med Hyg 85:124–131.

44. Dunn BE, Monson TP, Dumler JS, Morris CC, Westbrook AB, Duncan JL, Dawson JE, Sims KG, Anderson BE. 1992. Identification of Ehrlichia chaffeensis morulae in cerebrospinal fluid mononuclear cells. J Clin Microbiol 30:2207–2210.

45. Lyons JL, Schaefer PW, Cho TA, Azar MM. 2017. Case 34-2017. A 76-Year-Old Man with Fever, Weight Loss, and Weakness. N Engl J Med 377:1878–1886.

46. Huang C, Slater B, Rudd R, Parchuri N, Hull R, Dupuis M, Hindenburg A. 2002. First Isolation of West Nile virus from a patient with encephalitis in the United States. Emerg Infect Dis 8:1367–1371.

47. Morjaria S, Arguello E, Taur Y, Sepkowitz K, Hatzoglou V, Nemade A, Rosenblum M, Cavalcanti MS, Palomba ML, Kaltsas A. 2015. West Nile Virus Central Nervous System Infection in Patients Treated With Rituximab: Implications for Diagnosis and Prognosis, With a Review of Literature. Open Forum Infectious Diseases.

48. Levi ME, Quan D, Ho JT, Kleinschmidt-Demasters BK, Tyler KL, Grazia TJ. 2010. Impact of rituximab-associated B-cell defects on West Nile virus meningoencephalitis in solid organ transplant recipients. Clin Transplant 24:223–228.

49. Torres SD, Jia DT, Schorr EM, Park BL, Boubour A, Boehme A, Ankam JV, Gofshteyn JS, Tyshkov C, Green DA, Vargas W, Zucker J, Yeshokumar AK, Thakur KT. 2020. Central nervous system (CNS) enterovirus infections: A single center retrospective study on clinical features, diagnostic studies, and outcome. J Neurovirol 26:14–22.

50. Simner PJ, Miller HB, Breitwieser FP, Pinilla Monsalve G, Pardo CA, Salzberg SL, Sears CL, Thomas DL, Eberhart CG, Carroll KC. 2018. Development and Optimization of Metagenomic Next-Generation Sequencing Methods for Cerebrospinal Fluid Diagnostics. J Clin Microbiol 56.

51. Greninger AL. 2018. The challenge of diagnostic metagenomics. Expert Review of Molecular Diagnostics.

52. Hogan CA, Yang S, Garner OB, Green DA, Gomez CA, Dien Bard J, Pinsky BA, Banaei N. 2020. Clinical Impact of Metagenomic Next-Generation Sequencing of Plasma Cell-Free DNA for the Diagnosis of Infectious Diseases: A Multicenter Retrospective Cohort Study. Clin Infect Dis https://doi.org/10.1093/cid/ciaa035.

